# Nucleotide excision repair deficiency is a targetable therapeutic vulnerability in clear cell renal cell carcinoma

**DOI:** 10.1101/2023.02.07.527498

**Authors:** Aurel Prosz, Haohui Duan, Viktoria Tisza, Pranshu Sahgal, Sabine Topka, Gregory T. Klus, Judit Börcsök, Zsofia Sztupinszki, Timothy Hanlon, Miklos Diossy, Laura Vizkeleti, Dag Rune Stormoen, Istvan Csabai, Helle Pappot, Joseph Vijai, Kenneth Offit, Thomas Ried, Nilay Sethi, Kent W. Mouw, Sandor Spisak, Shailja Pathania, Zoltan Szallasi

## Abstract

**Purpose:** Due to a demonstrated lack of DNA repair deficiencies, clear cell renal cell carcinoma (ccRCC) has not benefitted from targeted synthetic lethality-based therapies. We investigated whether nucleotide excision repair (NER) deficiency is present in an identifiable subset of ccRCC cases that would render those tumors sensitive to therapy targeting this specific DNA repair pathway aberration.

**Experimental Design:** We used functional assays that detect UV-induced 6-4 pyrimidine-pyrimidone photoproducts to quantify NER deficiency in ccRCC cell lines. We also measured sensitivity to irofulven, an experimental cancer therapeutic agent that specifically targets cells with inactivated transcription-coupled nucleotide excision repair (TC-NER). In order to detect NER deficiency in clinical biopsies, we assessed whole exome sequencing data for the presence of an NER deficiency associated mutational signature previously identified in ERCC2 mutant bladder cancer.

**Results:** Functional assays showed NER deficiency in ccRCC cells. Irofulven sensitivity increased in some cell lines. Prostaglandin reductase 1 (PTGR1), which activates irofulven, was also associated with this sensitivity. Next generation sequencing data of the cell lines showed NER deficiency-associated mutational signatures. A significant subset of ccRCC patients had the same signature and high PTGR1 expression.

**Conclusions:** ccRCC cell line based analysis showed that NER deficiency is likely present in this cancer type. Approximately 10% of ccRCC patients in the TCGA cohort showed mutational signatures consistent with *ERCC2* inactivation associated NER deficiency and also substantial levels of *PTGR1* expression. These patients may be responsive to irofulven, a previously abandoned anticancer agent that has minimal activity in NER-proficient cells.

**Translational relevance:** DNA repair deficiencies can be therapeutically targeted by synthetic lethal-based strategies in cancer. However, clear cell renal cell carcinoma (ccRCC) has not benefitted from this therapeutic approach due to a lack of evidence for the presence of specific DNA repair pathway deficiencies. Here, we demonstrate that ccRCC harbors a therapeutically targetable DNA repair pathway aberration, nucleotide excision repair (NER) deficiency. ccRCC cell lines displayed robust signs of NER deficiency as determined by functional assays and some of these cell lines also displayed NER deficiency induced mutational signatures. These cell lines are also sensitive to irofulven, an abandoned anticancer agent that creates DNA lesions which can only be repaired by the NER pathway. We estimate that up to 10% of ccRCC cases may respond to NER-directed therapy with irofulven based on NER deficiency associated mutational signatures and PTGR1 expression levels, which is an enzyme required to activate irofulven.

## Introduction

Synthetic lethality driven therapy has become a successful treatment approach in the context of PARP inhibitor-based therapy for homologous recombination (HR) deficient ovarian, breast, prostate and pancreatic cancer. However, patients with clear cell renal cell carcinoma (ccRCC) have not benefitted from this treatment strategy thus far due to the absence of identifiable HR deficient cases. ccRCC cases almost never harbor inactivating mutations coupled with loss of heterozygosity (LOH) in the key HR genes (*BRCA1, BRCA2, RAD51* etc.). Furthermore, ccRCC cases rarely display DNA scarring signatures associated with HR deficiency (1). Therefore, it is likely that patients with ccRCC will not show sensitivity to PARP inhibitors via the mechanisms that confer sensitivity in ovarian or breast cancer. It was proposed recently that other genetic events, such as the inactivation of *PBRM1*, often observed in ccRCC, may confer PARP inhibitor sensitivity in this disease (2). However, the clinical relevance of this observation remains to be determined.

Nucleotide excision repair (NER) deficiency is another therapeutically targetable DNA repair deficiency in cancer. It is a highly conserved DNA repair pathway that recognizes and repairs bulky intrastrand DNA adducts (3). NER is initiated through two separate mechanisms of lesion recognition: transcription-coupled repair (TC-NER) is activated by RNA polymerase stalling at lesions, while global genome repair (GG-NER) is able to recognize distorted DNA structures throughout the genome. TC-NER and GG-NER converge on a common NER pathway that excises and replaces the damaged DNA strand in an error-free manner.

It has been known for decades that inactivation of NER activity in experimental models leads to increased cisplatin sensitivity. This is believed to be primarily driven by the ability of the NER pathway to remove platinum-induced DNA crosslinks. However, establishing a causative link between NER deficiency and platinum sensitivity in the clinic proved to be difficult due to the lack of diagnostic tools that detect NER deficiency in clinical biopsies. Recently, it was shown that mutations in the NER helicase gene *ERCC2* detected in urothelial carcinoma of the bladder cause NER deficiency in cell line models and that *ERCC2* mutations are associated with platinum sensitivity in some bladder cancer clinical cohorts (4,5). Thus, preliminary evidence for NER deficiency and associated platinum sensitivity was established in at least one solid tumor type. Indirect evidence for the presence of NER deficiency is also presented in other solid tumor types as well, such as breast cancer (6). We previously reported increased risk for breast cancer due to recurrent *ERCC3* variant and demonstrated lower cell survival in mutant mammary epithelial cell line (HMLE), when exposed to IlludinS, a DNA damaging sesquiterpene (7).

Here we provide experimental evidence for the presence of NER deficiency in ccRCC cell lines. We also demonstrate that the specific mutational signatures associated with *ERCC2* inactivation in bladder cancer are also present in a subset of ccRCC cases. Finally, we show that the mutational signature of NER deficiency detected in ccRCC cell lines is associated with increased sensitivity to irofulven, an experimental therapeutic agent with synthetic lethal activity in NER deficient cells.

## Materials and Methods

### Cell lines and reagents

Cell lines 786O, 769P, A498 were purchased from ATCC^®^. SLR26, CAKI1, ACHN and RXF393 were kindly supplied by the Kaelin laboratory (Dana Farber Cancer Institute). Cell lines were grown in RPMI 1640 (Gibco) supplemented with 10% FBS (Gibco), incubated at 37°C in 5% CO2, and regularly tested for Mycoplasma spp. contamination.

The NCI-H460 cell line was purchased from ATCC. The Alt-R™ CRISPR-Cas9 System (IDT Technologies) was used to delete *ERCC4*. Cas9 nuclease was purchased from Horizon Discovery. The crRNA was annealed with ATTO™ 550-tracrRNA, and ribonucleoparticles (RNPs) were then assembled by adding Cas9. RNPs were delivered into cells using electroporation-based nucleofection (Lonza system). Flow cytometry was utilized to sort ATTO-550 positive single cells 24 hours following nucleofection. Next, single cells were expanded and clonal populations were screened by immunoblot to identify clones with complete loss of expression of the ERCC4 protein.

### *In vitro* drug sensitivity assays

Exponentially growing cell lines were seeded in 96-well plates (3000 cells/well) and incubated for 24 hrs to facilitate cell attachment. Identical cell numbers of seeded parallel isogenic lines were verified by the Celigo Imaging Cytometer after attachment. Cells were exposed to Irofulven (Cayman Chemicals) for 72 hrs, and cell growth was determined by the addition of PrestoBlue (Invitrogen) and incubated for 2.5 hrs. Cell viability was determined by using the BioTek plate reader system. Fluorescence was recorded at 560 nm/590 nm, and values were calculated based on the fluorescence intensity. IC50 values were determined by using the AAT Bioquest IC50 calculator tool. P-values were calculated using student’s t-test. P-values <0.05 were considered statistically significant.

### PTGR1 knockdown

An siRNA against PTGR1 (ON-TARGETplus; Dharmacon), shown to induce >90% reduction of PTGR1 transcript levels over 48-72 hours, or an Alexa Flour non-targeting control siRNA were transfected at 25nM into the HMLE cell line using Lipofectamine RNAiMAX (Thermo Scientific). Cells were seeded at 3000 cells per well into a 96-well plate during reverse transfection. Following 24 hours, the cells were treated with either vehicle (0.01 % EtOH) or irofulven at 300 nM and 600 nM doses. Cell viability was measured after 72 hours using the CellTiterGlo reagent (Promega).

### Immunoblotting

Freshly harvested cells were lysed in RIPA buffer. Protein concentrations were determined by Pierce BCA^TM^ Protein Assay Kit (Pierce). Proteins were separated via Mini Protean TGX stain free gel 4-15% (BioRad) and transferred to polyvinylidene difluoride membrane by using iBlot 2 PVDF Regular Stacks (Invitrogen) and iBlot system transfer system (LifeTechnologies). Membranes were blocked in 5% BSA solution (Sigma). Primary antibodies were diluted following the manufacturer’s instructions: anti-beta Actin, [AC15] (HRP-conjugated) ab 49900, Abcam (1:25000) and antiPTGR1 [EPR13451-10], ab181131, Abcam (1:1000). Signals were developed using Clarity Western ECL Substrate (BioRad) and Image Quant LAS4000 System (GEHealthCare).

### NER Assay

Removal of 6-4 pyrimidine-pyrimidone photoproducts (6-4PP) as a function of NER was quantified using an immunofluorescent assay. Cells on coverslips were fixed in cold methanol for 10 minutes on ice, and triton was extracted (0.5% Triton X-100 in PBS) for 4 minutes at room temperature. The coverslips were then incubated at 37 °C for 15 minutes in 2M HCL in PBS. After washing twice with PBS, once with 1% BSA/PBS, once with PBS, cells were incubated with 6-4PP primary antibody (NM-DND-002, 1:2000) for 45 minutes at 37°C followed by incubation with secondary antibody for 30 minutes at 37°C. Coverslips were then washed twice with PBS and mounted using DAPI.

### Patients and cell lines

This study evaluated 389 whole exome sequenced (WES) pretreatment samples of RCC patients from the TCGA-KIRC cohort. The normal, tumor bam and vcf files were retrieved from the TCGA data portal (https://portal.gdc.cancer.gov/) for the analysis. From the TCGA data portal the vcf files for the somatic mutations from the MuTect2 pipeline were used.

Variants were collected from the DepMap portal (https://depmap.org/portal/download/) for the cancer cell line samples (DepMap version 22Q2).

### Mutation calling and filtering

The application of the MuTect2 default filters (FILTER == “PASS”) for filtering the called mutations ensured the high accuracy of germline and somatic changes reported. Utilizing additional stringent filters on somatic samples provided the high accuracy of reported variants: TLOD ≥ 6 and NLOD ≥ 3, NORMAL.DEPTH ≥ 15 and TUMOR.DEPTH ≥ 20, TUMOR.ALT ≥ 5 and NORMAL.ALT = 0 and TUMOR.AF ≥ 0.05. Additionally, samples with less than a total of 50 variants were removed, since mutational signature extraction is less reliable when the number of mutations is fewer than 50.

After applying these filters and keeping only one sample per patient (by removing the samples with whole genome amplification) and removing the FFPE samples and samples indicated having MSI (Microsatellite Instability) using the MANTIS tool (8) 289 samples were further analyzed.

Intervar (version 2.0.2) was utilized to classify the variants as “Benign,” “Likely Benign,” “Uncertain Significance,” “Likely Pathogenic,” and “Pathogenic.” Deleterious mutations were defined for exonic SNVs with “Pathogenic” or “Likely Pathogenic” labels, nonsense SNV-s and indels with “Pathogenic” or “Likely Pathogenic” labels. All the ERCC gene family mutants represented in the figures are deleterious mutations.

For genotyping of the cell line samples, variants were defined as deleterious if the column “isDeleterious” was indicated as “True” in the CCLE.mutations.csv data file.

### Mutational signatures

Using techniques based on non-negative matrix factorization, Alexandrov et al. (9) described single base substitutions (SBS) signatures, doublet base substitution (DBS) signatures and small insertion and deletion (ID) signatures. In this study we calculated the number of ID8 signatures since we previously found this signature most significantly associated with NER deficiency (10). The identified matrix of ID signatures was downloaded from https://www.synapse.org/#!Synapse:syn12025148. ID mutations in each sample were classified into 83-dimensional indel catalog using the ICAMS R package (11). The resulting matrices were used in a non-negative least-squares problem to estimate the matrix of exposures to mutational processes.

The ID8 signature extraction was performed the same way on the patient and cancer cell line samples.

### RNA expression analysis

RNA expression data were downloaded from the TCGA data portal (https://portal.gdc.cancer.gov/) for the patient samples, and The Fragments Per Kilobase of Transcript per Million Mapped Reads (FPKM) technique was used to normalize the data, and the data were log2-transformed using a pseudo-count thereafter.

For the cancer cell line samples, the RNA expression data were obtained from the DepMap portal (https://depmap.org/portal/) and the TPM-normalized data were log2-transformed using a pseudo-count. For comparison with the TCGA-KIRC PTGR1 FPKM values, cell-line expression data in FPKM were downloaded from the CellMiner website (https://discover.nci.nih.gov/cellminer/).

### Code availability

There are no restrictions to accessing the custom code used for the analyses presented in this study. Information is available from the authors on request.

## Results

### A subset of ccRCC cell lines are highly sensitive to irofulven

Cancer cells with defective transcription coupled repair show approximately 100-fold increased sensitivity to irofulven (12). Interestingly, drug sensitivity experiments from the NCI60 drug screening program reported that RXF393, a kidney cancer cell line, showed high sensitivity to irofulven (https://dtp.cancer.gov/services/nci60data/colordoseresponse/jpg/683863). Recently it was also reported that the ccRCC cell lines A498 and RXF393, also show significant sensitivity (IC50~20nM) to a recently developed analog of irofulven (13). We expanded these experiments to include a panel of seven kidney cancer cell lines (Figure 1). A498 had an IC50 of 91nM and RXF393 had an IC50 of 153nM, well below the estimated plasma concentration of 400nM irofulven that was achieved in patients without significant dose limiting toxicities (14). These IC50 values also place these cell lines among the most sensitive to irofulven and its analog among a wide variety of solid cancer types (13).

**Figure 1:**
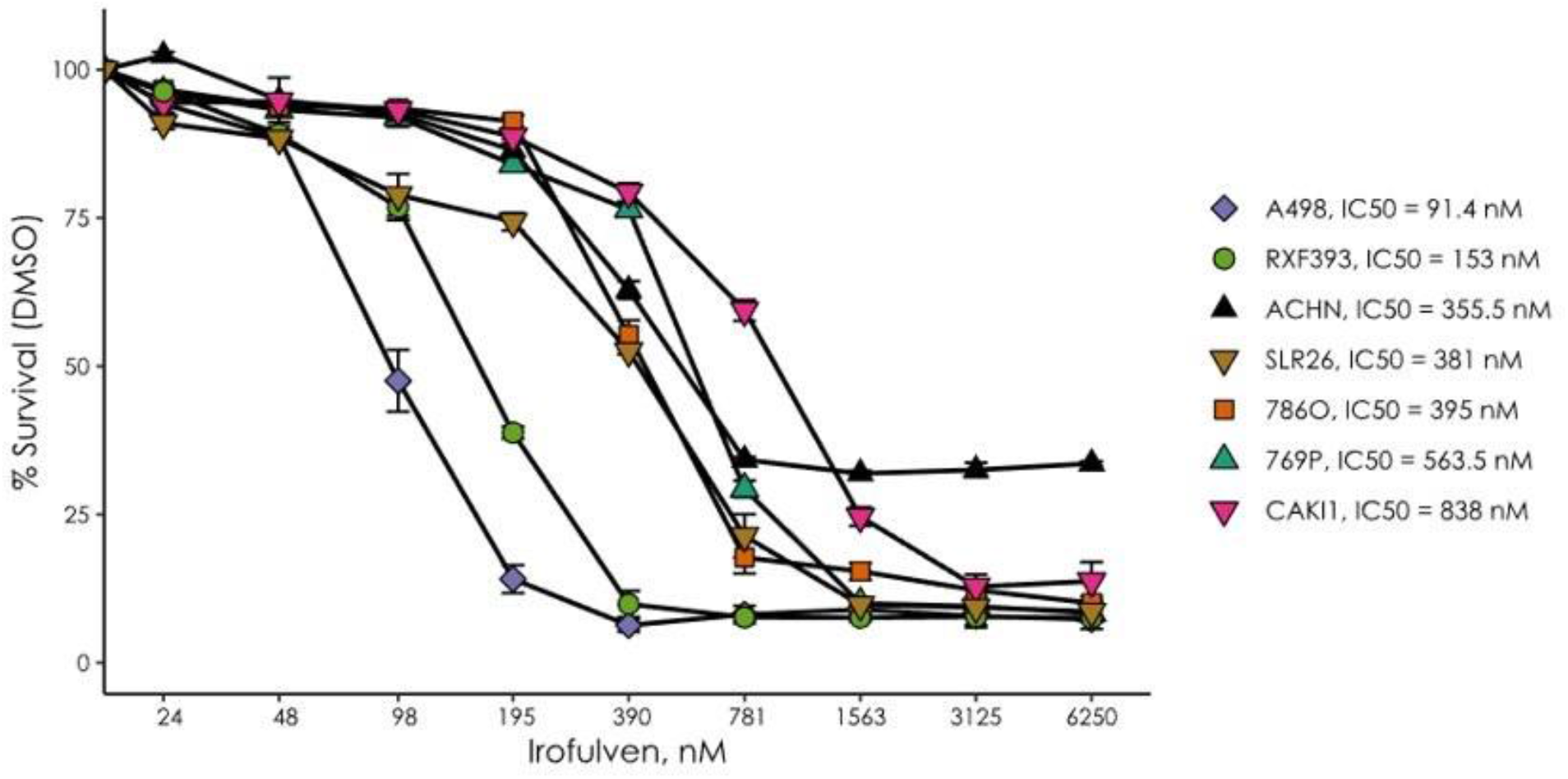
Kidney cancer cell lines show various degrees of sensitivity to irofulven. In vitro cell viability assays indicating some cell lines having an effective IC50 around 100 nM. Kidney cancer cell lines were incubated with various concentrations of irofulven for 72 hrs as indicated and cell viability was measured using PresoBlueTM reagent. The error bars represent the mean plus and minus the standard error.

### Clear cell renal cell carcinoma cell lines show various degrees of nucleotide excision repair deficiency by functional assays

One of the prerequisites of irofulven sensitivity is defective nucleotide excision repair (12). We performed a functional assay of NER wherein NER efficiency was determined by monitoring the repair of UV-induced 6-4PP photoproducts in the clear cell renal carcinoma cell lines. We analyzed the NER efficiency in ccRCC cell lines with a functional assay of NER as described in the clear cell renal carcinoma cell lines with high sensitivity (A498, RXF393) and low sensitivity (786O and 769P) to irofulven, in the non-malignant immortalized HK-2 kidney epithelial cell line. As mentioned above, this assay monitors the cells’ ability to remove UV-induced 6-4 pyrimidine-pyrimidone photoproducts (6-4PP). 6-4PPs can be removed by both GGR (global genome repair) and TCR (transcription coupled repair) pathways of NER and their removal is closely correlated with NER efficiency (15). Using this assay, we found that surprisingly all five kidney epithelial cell lines, including the non-malignant HK2 cells, showed NER deficiency to varying degrees. In contrast, the control cell line (the NER proficient H460 cell line) was NER proficient and efficiently removed the 6-4PP photoproducts by 7 hours post UV irradiation (Figure 2). As a positive control for NER deficiency, we used the H460 cell line in which *ERCC4*, a key NER gene, was deleted using CRISPR-Cas9 methodology. As expected for an NER deficient line, H460 ERCC4 KO line shows no repair of 6-4PP by 7hrs post UV. The RXF393 cell line had a level of NER deficiency similar to that detected in a cell line with a complete loss of *ERCC4*.

**Figure 2:**
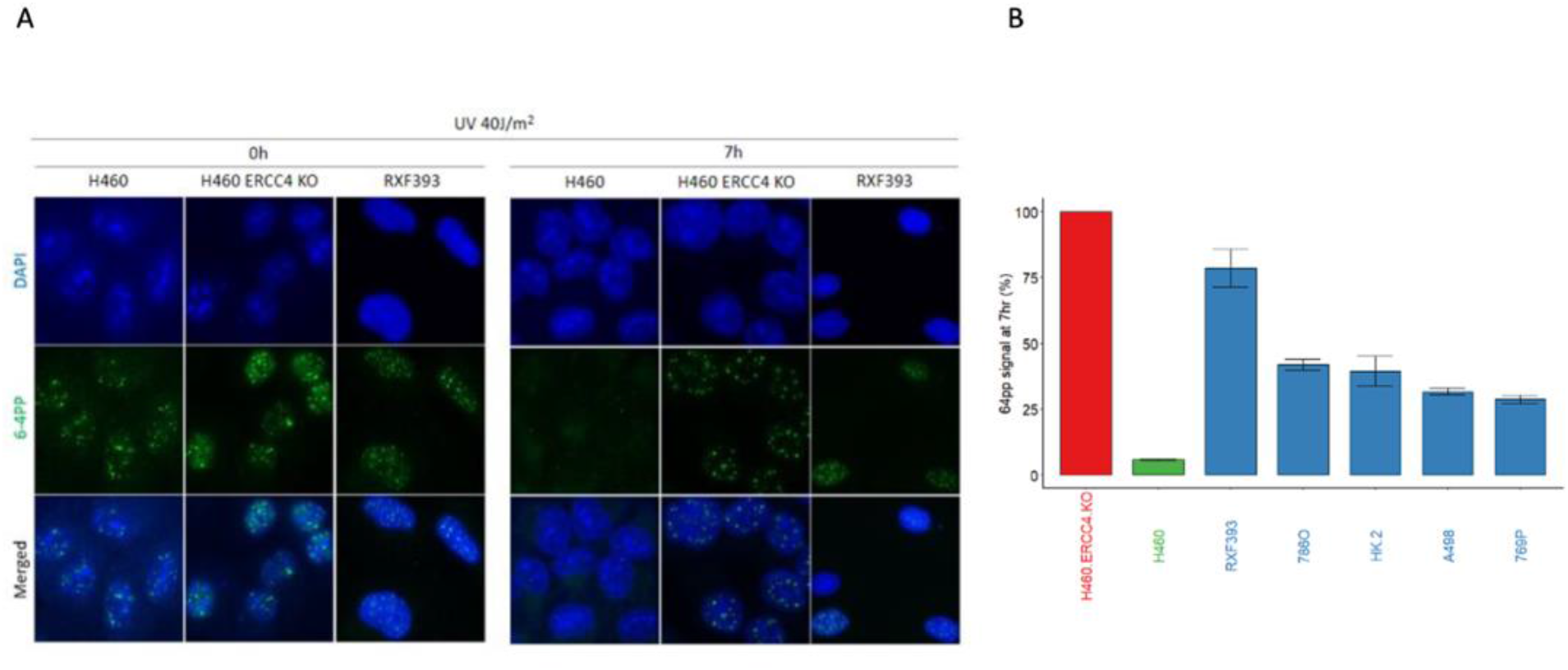
Kidney cancer cell lines show various degrees of nucleotide excision repair deficiency by a functional assay monitoring the cells’ ability to remove 6-4-photoproducts. A, Cells were irradiated by UV and 6-4-photoproducts were detected as described at 0 and 7 hours. NER activity is expressed by the percent of 6-4-photoproducts removed by 7 hours after UV irradiation. The H460 cell line and its engineered, ERCC4 deficient derivative was used as positive and negative controls. B, On the barplot the mean of the measurements is shown 7 hours after the UV irradiation, normalized by the signal at measured at 0 hours. The two whiskers represent the mean plus and minus the standard error.

### PTGR1, a functionally validated prerequisite of irofulven sensitivity, is expressed in several kidney cancer cell lines

Irofulven acts as a prodrug, and overexpressing the metabolic activator prostaglandin reductase 1 (*PTGR1*) increases its efficacy (16). Here we provide direct functional evidence that the presence of *PTGR1* is a key determinant of drug response by demonstrating that suppression of *PTGR1* expression in an otherwise irofulven sensitive, NER deficient cell line renders those cells irofulven resistant. A heterozygous truncating mutation in *ERCC3* (p.R109X) was previously introduced by CRISPR editing into the HMLE cell line (7,17). The mutation rendered these cells sensitive to irofulven. We depleted *PTGR1* in these cells with siRNA and found that depletion of *PTGR1* rendered those cells resistant to irofulven (Figure 3A).

**Figure 3:**
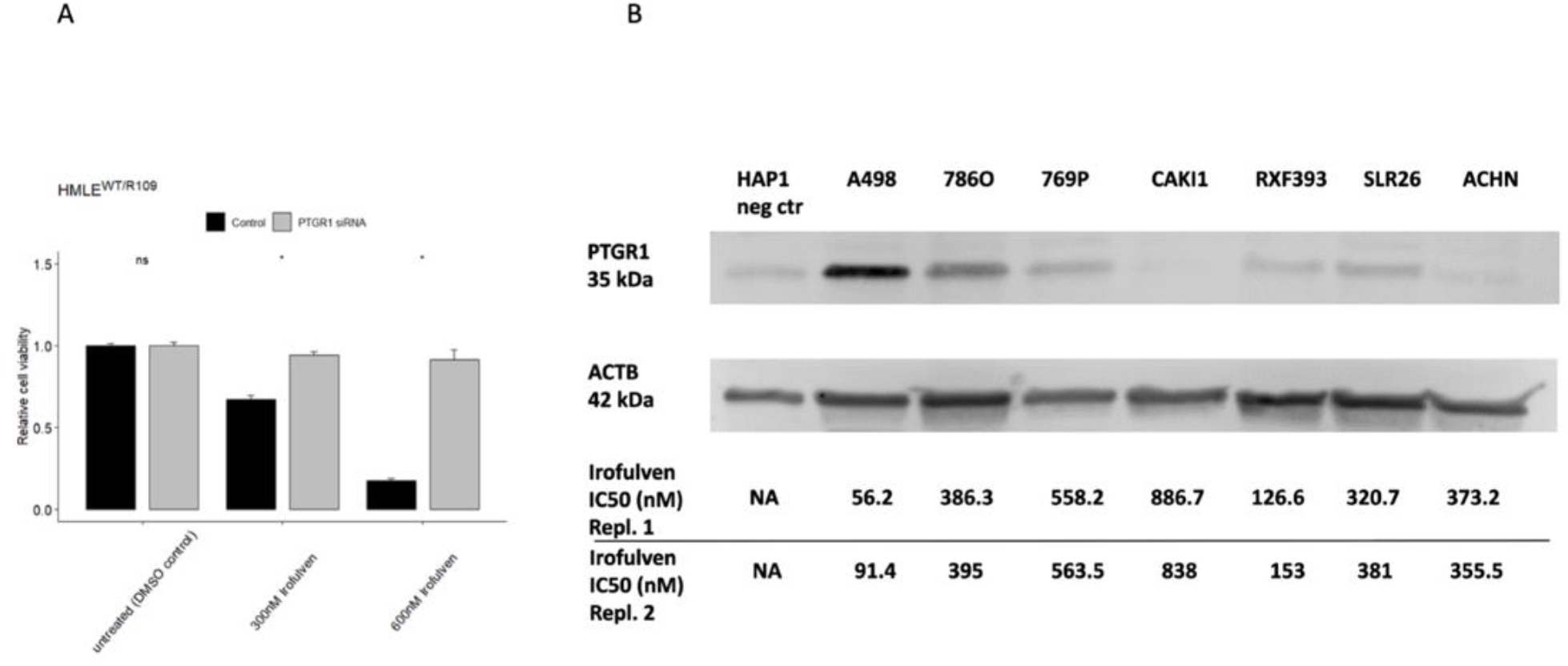
PTGR1, a functionally validated prerequisite of irofulven sensitivity, is expressed in several kidney cancer cell lines. **A)** R109X mutant version of ERCC3 was introduced into the HMLE cell line rendering these cells sensitive to irofulven as previously described (17). Suppressing PTGR1 by siRNA has restored resistance to irofulven. **B)** PTGR1 protein expression levels were determined by Western blot analysis in the various kidney cancer cell lines. Irofulven sensitivity of the individual cell lines is indicated at the bottom.

Since *PTGR1* expression is one of the possible determinants of irofulven sensitivity, we quantified *PTGR1* expression by Western blot analysis in the above listed cell line panel. With the exception of two irofulven resistant cell lines (CAKI1 and ACHN), all other kidney cancer cell lines expressed *PTGR1* and a trend could be observed between *PTGR1* expression levels and irofulven sensitivity, although the limited number of cell lines did not allow establishing a statistically significant correlation (Supplementary Figure 1). It is notable, however, that one of the two most irofulven sensitive cell lines had the highest expression of *PTGR1* (A498 in Figure 2B) and the other highly sensitive line showed the highest level of NER deficiency by the functional assay (RXF393 on Figure 1) (Supplementary table 1).

### NER deficiency of ccRCC cell lines is associated with a NER deficiency specific mutational signature

NER deficiency can be functionally assessed as described above, but these methods cannot currently be applied to clinical biopsies. We recently identified a set of mutational signatures strongly associated with *ERCC2* inactivating mutations (10). Most prominent of these NER-related signatures is ID8, which is a mutational signature characterized by longer than 5 bp deletions with no or short 1-2 bp flanking microhomologies. We assessed whether the ID8 signature is present in the whole exome sequencing data of ccRCC cell lines. These cell lines display various levels of ID8 signature deletions but all four cell lines (A498, RXF393, 786O, 769P) that showed NER deficiency by the functional assay also had a high level of ID8 deletions (Figure 4). Conversely, the cell lines we used in our functional assay as NER proficient controls (H460 as well as HCT116 and HeLa) had a low number of ID8 deletions. These results suggest that the NER deficiency-associated mutational signature ID8 may be indicative of NER deficiency in kidney cancer cells.

**Figure 4:**
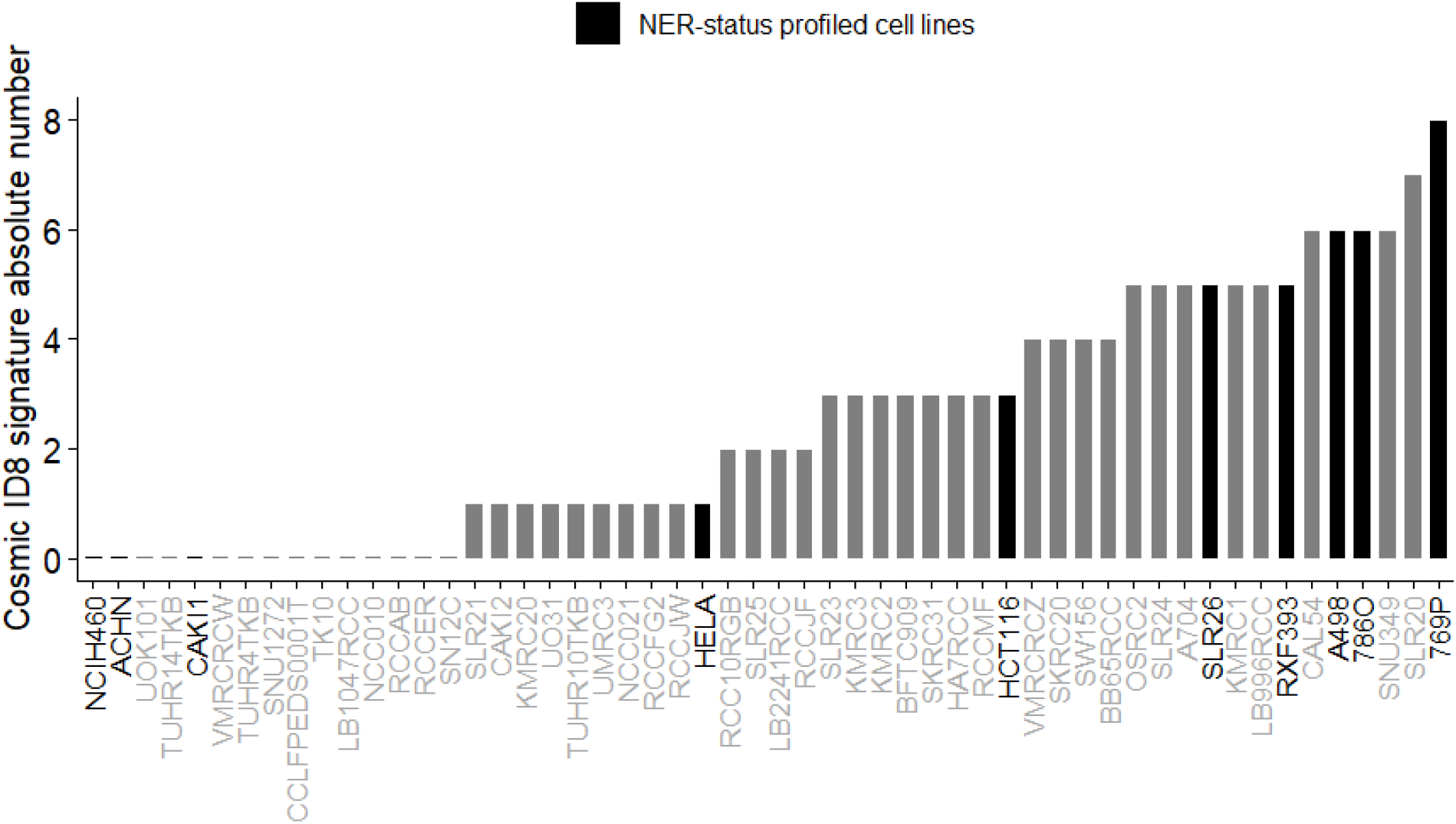
The distribution of the absolute number of Cosmic mutational signature ID8 in a selected panel of cancer cell lines. ID8 deletions were extracted from the whole exome sequencing data of a panel of ccRCC cell lines and two control cell lines with active NER function (HeLA and HCT116). The cell lines profiled in our NER functional assay experiments are highlighted.

### A subset of ccRCC clinical cases display the mutational signature of NER deficiency and *PTGR1* expression

We have shown previously that NER deficiency associated mutational events are enriched in actively transcribed genomics regions, therefore whole exome sequencing data can be used to detect likely NER deficient cases (10). 289 cases of the TCGA ccRCC cohort passed our quality control for further analysis (see methods). We identified four cases predicted deleterious mutations in *ERCC6*, three cases with predicted deleterious mutations in *ERCC2*, one case with a predicted deleterious in *ERCC3*, and one case with multiple NER gene mutations (*ERCC2, ERCC3* and *ERCC6*). These cases with NER gene mutations showed a statistically significant association with higher ID8 events (Figure 5A, Fisher’s p = 0.00018). We previously established that more than five ID8 deletions detected in WES analysis indicates the likely presence of NER deficiency in bladder cancer (10). We used the same threshold in kidney cancer and found that 43 out of 289 cases (~15%) had ID8 deletion numbers consistent with NER deficiency.

**Figure 5:**
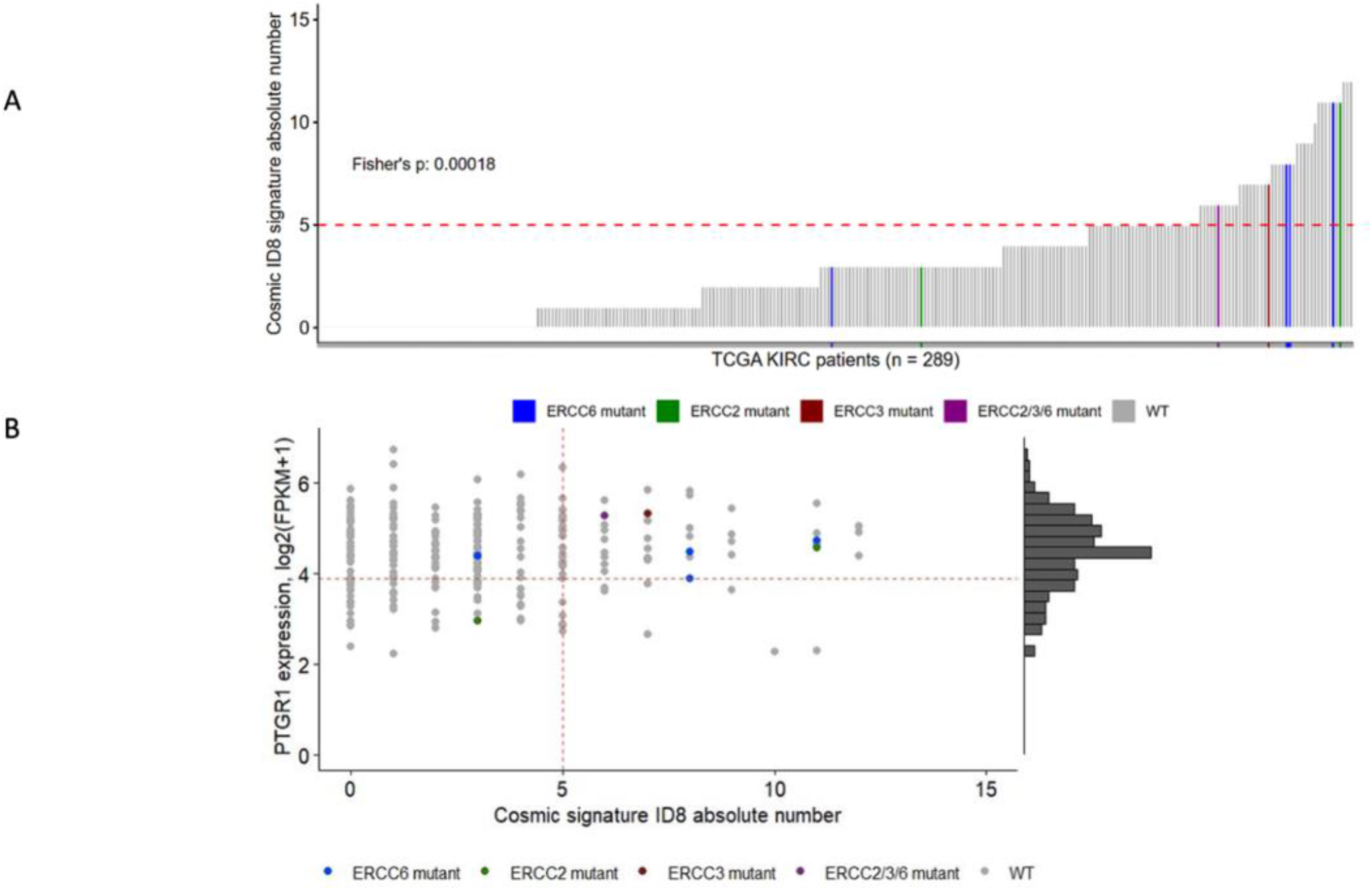
A significant portion of patients from the TCGA-ccRCC cohort have high Cosmic mutational signature ID8 absolute numbers and high PTGR1 expression. A) Ordered Cosmic mutational signature ID8 absolute number distribution in the TCGA-ccRCC cohort. Cut-off value of >5 of ERCC2 mutation induced NER deficiency was previously defined in bladder cancer cohorts (dashed-line) (10). Cases with ERCC2, ERCC3 or ERCC6 mutations are highlighted, where the forward slash (/) symbol means multiple ERCC gene family mutations in the same sample. Fisher exact test was performed between the samples below and above the cutoff value if they harbor any ERCC gene family mutations. B) Joint distribution of patients from the TGCA-ccRCC cohort regarding to the Cosmic mutational signature ID8 absolute number and PTGR1 expression. Samples with ERCC gene family mutations and high Cosmic mutational signature ID8 tend to also have high PTGR1 expression. The cut-off value for the PTGR1 expression was determined on the renal cancer cell-lines.

We also estimated the expression levels of *PTGR1* using TCGA RNAseq data and compared those to the *PTGR1* expression levels detected in the ccRCC cell lines. 36 of the 43 cases with >5 ID8 deletions had the same or higher level *PTGR1* expression as the A498 cell line that had the highest level of PTGR1 expression at the protein level and also had a high level of irofulven sensitivity (Figure 3B). Such cases likely have the sufficient level of *PTGR1* activity to activate irofulven.

Considering these criteria 36 of the total number of 389 TCGA cases (~9%) indicated the presence of both NER deficiency and significant *PTGR1* expression levels thus defining the proportion of clear cell renal carcinoma cases that may respond to irofulven therapy (Figure 5B).

## Discussion

Tumor DNA repair deficiency can be therapeutically targeted by synthetic lethal–based strategies. The success of the synthetic lethal approach is dependent on the identification of the relevant DNA repair pathway deficiency in clinical tumor specimens and the availability of a therapeutic agent that can specifically target such DNA repair–deficient cells. ccRCC in general has not benefitted from this therapeutic strategy because the presence of specific DNA repair pathway deficiencies has not been demonstrated in this tumor type. Here we show that NER deficiency can be detected in several ccRCC cell line models by functional assays and that a subset of clinical ccRCC cases have mutational features consistent with NER deficiency.

Since currently there are no functional or IHC assays available to reliably identify NER deficiency from clinical specimens, we used a specific mutational signature (ID8) associated with *ERCC2* helicase inactivating mutations (10). We were particularly encouraged by the fact that ccRCC is one of the solid tumor types where the highest proportion of cases harbor this mutational signature and also that the frequency of ID8 deletions is also among the highest across the various solid tumor types (9). In our analysis, ID8 was present both in some of the ccRCC cell lines and patient biopsies at levels detected in *ERCC2* mutant bladder cancer cases. Furthermore, this mutational signature was also associated with either the presence of functional NER deficiency (cell lines) or inactivating mutations in NER genes (TCGA biopsies). This suggests that the *ERCC2* mutation associated mutational signature we previously described may also indicate the presence of NER deficiency in ccRCC. The ID8 signature may, however, be caused by other mechanisms as well. A rare somatic mutation of topoisomerase II alpha was previously described to be associated with this signature before (18). This may lead to an overestimation of truly NER deficient ccRCC cases.

Our interest in the diagnostic detection of NER deficiency in ccRCC was inspired by the remarkable sensitivity of some of the commonly used ccRCC cell lines to irofulven, which is a semisynthetic, DNA alkylating agent that is a derivative of the fungal sesquiterpene, illudin S (19). Cells with inactivated transcription couple repair (TCR) or NER show an approximately 100-fold increased cytotoxic activity relative to normal cells with active DNA repair (12). This suggests an exploitable therapeutic index for NER deficient cases. However, although well-tolerated, irofulven showed only modest clinical benefit as a single agent in phase I/II clinical trials across a variety of tumor types (20–22) including a phase II trial for advanced renal cell carcinoma (23). The failure of irofulven to show clinical benefit in this limited set of thirteen renal cell carcinoma patients may be due to the fact that patients were not selected according to the two criteria for irofulven activity: NER deficiency and the expression of *PTGR1*. According to these criteria, we estimate that approximately one in ten ccRCC patients may respond to irofulven. Therefore, in the case of thirteen unselected patients, it is not surprising that no NER-deficient cases were included. In a basket trial of irofulven/cisplatin combination therapy, four ccRCC patients were included and one experienced a significant partial response (24).

NER deficient cells are particularly sensitive to platinum treatment (4). Early disappointing phase II clinical trials with single agent platinum or platinum combination therapies suggested limited use for this form of treatment in ccRCC (25,26). In these trials a 5% objective response rate was observed with limited cohort sizes (~20 patients each). The low objective response precluded the development of larger, biomarker directed trials for platinum. Therefore, currently we do not know whether the likely NER deficient subset of patients would benefit from platinum-based therapy. It is notable, however, that one of the NER deficient, irofulven sensitive cell line in our analysis, RXF393, has been reported to be as sensitive to platinum treatment as the NER deficient breast cancer cell line, MDA-MB468 (6) or the homologous recombination deficient breast cancer cell line, MDA-MB436 (Genomics of Drug Sensitivity in Cancer, www.cancerrxgene.org).

*PTGR1*, the enzymatic activator of irofulven, is a NADPH-dependent alkenal/one oxidoreductase with high expression levels in the kidney, the tissue from where it was originally cloned (27). The significant expression of this enzyme in the majority of kidney cancer cases is perhaps a result of the retention of a key pathway for kidney metabolism of leukotrienes. Since *PTGR1* is not expressed in every cell type, (e.g., there is a notable complete lack of expression in white blood cells (27)), several normal tissues are not affected by the toxicities of irofulven treatment due to lack of enzymatic activation of the drug. This significantly contributes to the good tolerability, including its lack of hematological toxicity (20), while potentially retaining the majority of NER deficient kidney cancer cases as a potential therapeutic target.

One of the unexpected results of our experiments was the fact that all kidney epithelium cell lines, including a non-malignant cell line, showed signs of NER deficiency. It was shown before that hypoxia-inducible factor-1α regulates the expression of nucleotide excision repair proteins in keratinocytes (28). Therefore, it is possible that the NER deficiency we detected in several kidney epithelial cell lines may in fact be the result of culturing those cells under conditions, in this case normoxia, that would lead to the inactivation of NER. This would also suggest that under hypoxic conditions, when the risk of the various forms of DNA damage is increased, NER would be reactivated. In theory, if the underlying molecular mechanisms can be identified, then inactivating NER by such an oxygen sensing mechanism could also sensitize a wider range of ccRCC cases to NER deficiency targeted therapy.

Taken together, we estimate that about 10% of ccRCC cases may be responsive to irofulven therapy and a biomarker directed clinical trial could identify this population.

## Authors’ Contributions

A. Prosz: Conceptualization, formal analysis, visualization, methodology, writing–original draft, writing–review and editing. H. Duan: Conceptualization, formal analysis, visualization, methodology, writing–original draft, writing–review and editing. V. Tisza: Conceptualization, formal analysis, visualization, methodology, writing–original draft, writing–review and editing. P Sahgal: Conceptualization, investigation, writing–review and editing. S. Topka: Conceptualization, investigation, writing–review and editing. G.T. Klus: Conceptualization, investigation, writing–review and editing. J. Borcsok: Conceptualization, formal analysis, methodology, writing–review and editing. Z. Sztupinszki: Conceptualization, formal analysis, methodology, writing–review and editing. T. Hanlon: Conceptualization, investigation, writing– review and editing. M Diossy: Conceptualization, formal analysis, methodology, writing–review and editing. L. Vizkeleti: Conceptualization, investigation, writing–review and editing. D.R. Stormoen: Writing–review and editing. I. Csabai: Writing–review and editing. H. Pappot:

Writing–review and editing. J.Vijai and K. Offit:: Funding acquisition, writing–review and editing. T. Ried: Funding acquisition, writing–review and editing N. Sethi: Funding acquisition, writing–review and editing. K.W. Mouw: Conceptualization, supervision, funding acquisition, investigation, writing–original draft, writing–review and editing. S. Spisak: Conceptualization, supervision, funding acquisition, writing–original draft, writing–review and editing. S. Pathania: Conceptualization, supervision, funding acquisition, writing–original draft, writing–review and editing Z. Szallasi: Conceptualization, supervision, funding acquisition, writing–original draft, writing–review and editing.

## Acknowledgments

The results shown here are partly based upon data generated by the TCGA Research Network: *http://cancergenome.nih.gov/*

## Ethical Statement

The authors are accountable for all aspects of the work in ensuring that questions related to the accuracy or integrity of any part of the work are appropriately investigated and resolved.

---

**Supplementary table 1:** PTGR1 expression, Irofulven IC50 and 6-4PP IF ratio at the 7^th^ hour for the NER-profiled cell lines.

| Cell-line | PTGR1 expression | IC50 (nM) | 6-4PP IF |
| --- | --- | --- | --- |
| A498 | +++ | 91.4 | 31.6 |
| 786O | ++ | 395 | 41.8 |
| 769P | + | 563.5 | 28.7 |
| CAKI1 | - | 838 | NA |
| RXF393 | + | 153 | 78.6 |
| SLR26 | + | 381 | NA |
| ACHN | - | 355.5 | NA |

**Supplementary figure 1:**
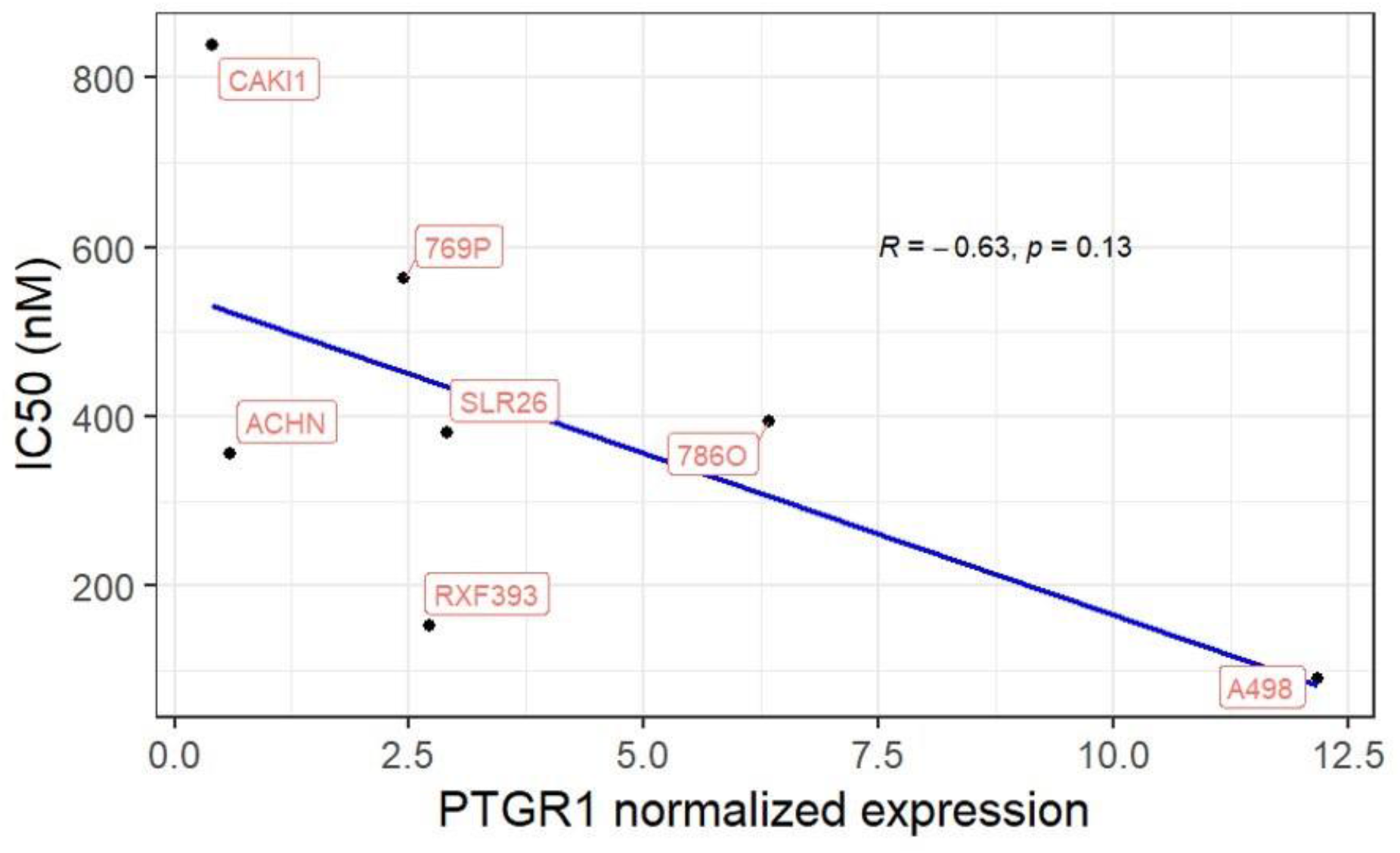
A negative trend can be observed between PTGR1 normalized expression and Irofulven IC50 values in renal cancer cell lines. The relationship between the expression level of PTGR1 determined by Western Blot and the effectiveness of Irofulven in preventing the proliferation of kidney cancer cells. Elevated PTGR1 expression correlates with enhanced medication potency (lower IC50 values).

## Notes

**Additional information**, <u>Financial support</u>: This work was supported by the Research and Technology Innovation Fund (KTIA_NAP_13-2014-0021 to Z.S.), Breast Cancer Research Foundation (BCRF-21-159 to Z.S.), the Novo Nordisk Foundation Interdisciplinary Synergy Programme Grant (NNF15OC0016584 to Z.S.), Kræftens Bekæmpelse (R281-A16566 to Z.S. and R340-A19380 to J.B.), Department of Defense through the Prostate Cancer Research Program (W81XWH-18-2-0056 to Z.S.), Det Frie Forskningsråd Sundhed og Sygdom (7016-00345B to Z.S.), the National Cancer Institute (R01CA272657 to K.W.M), and the Velux Foundation (00018310 to Zs.S. and J.B.). This work was also supported by a grant from The National Cancer Institute, R15 CA 235436-01 (S.P.). We acknowledge the support of the NIH core grant to MSKCC (P30 CA008748), the MSKCC bladder SPORE (CA221745), the breast cancer research foundation (BCRF) grant and the Kate and Robert Niehaus Foundation providing funding to the Robert and Kate Niehaus Center for Inherited Cancer Genomics at Memorial Sloan Kettering Cancer (KO). S.S. received funding from National Research Development and Innovation Office Hungary, under grant no. FK142835.

**<u>Conflict of interest:</u>** K.W.M - Consulting or Advisory Role: EMD Serono, Pfizer. Research Funding: Pfizer. Patents: Institutional patents filed on *ERCC2* mutations and chemotherapy response (KW.M, Z.S., J.B, Zs. Sz. and M.D.). JV, ST and KO are inventors on a patent application for use of Illudin class of alkylating agents in patients harboring mutations in the *ERCC3* gene (PCT/US2018/022588). D.R.S: Research Funding: Pfizer, EMD Serono.H.P.: Research funding from Pfizer and Merck Z.S: Research funding from Lantern Pharma Inc.

### Competing Interest Statement

K.W.M - Consulting or Advisory Role: EMD Serono, Pfizer. Research Funding: Pfizer. Patents: Institutional patents filed on ERCC2 mutations and chemotherapy response (KW.M, Z.S., J.B, Zs. Sz. and M.D.). JV, ST and KO are inventors on a patent application for use of Illudin class of alkylating agents in patients harboring mutations in the ERCC3 gene (PCT/US2018/022588). D.R.S: Research Funding: Pfizer, EMD Serono.H.P.: Research funding from Pfizer and Merck Z.S: Research funding from Lantern Pharma Inc.

